# Mapping the Location of Nucleosomes in Simian Virus 40 Chromatin Using the New England Biolabs FS Library Prep Kit

**DOI:** 10.1101/2020.03.27.011924

**Authors:** Barry Milavetz, Jacob Haugen, Kincaid Rowbotham

**Affiliations:** Department of Biomedical Sciences, University of North Dakota School of Medicine and Health Sciences, Grand Forks, 58202, USA

## Abstract

The location of nucleosomes in chromatin significantly impacts many biological processes including DNA replication, repair and gene expression. A number of techniques have been developed for mapping nucleosome locations in chromatin including MN-Seq (micrococcal nuclease digestion followed by next generation sequencing), ATAC-Seq (Tagamet chromatin fragmentation followed by next generation sequencing), and ChIP-Seq (chromatin immunoprecipitation and fragmentation followed by next generation sequencing). All of these techniques have been successfully used, but each with its own limitations. Recently, New England Biolabs has marketed a new kit, the NEBNext UltraII FS Library Prep kit, for preparing libraries for next generation sequencing from purified genomic DNA. This kit is based on a novel proprietary DNA fragmentation procedure which appears to cleave DNA that is not bound by proteins. Because DNA is fragmented directly in the FS kit, we tested whether the kit might also be useful for mapping the location of nucleosomes in chromatin. Using Simian Virus 40 (SV40) chromatin isolated at different times in an infection, we have compared nucleosome mapping using the NEB FS kit (FS-Seq) to MN-Seq, ATAC-Seq, and ChIP-Seq. Mapping nucleosomes using FS-Seq generated nucleosome profiles similar to those generated by ATAC-Seq and ChIP-Seq in regulatory regions of the SV40 genome. We conclude that FS-Seq is a simple, robust, cost-effective procedure for mapping nucleosomes in SV40 chromatin that should be useful for other forms of chromatin as well. We also present evidence that the FS kit may be useful for mapping the location of transcription factors in chromatin when sequencing reads between 75 and 99 base pairs in size are analyzed.

## Introduction

The organization of nucleosomes in chromatin is thought to play a role in epigenetic regulation by controlling the accessibility of regulatory DNA sequences in the chromatin to recognition by transcription and replication factors. For example, a change in nucleosome organization is associated with the activation of transcription and is typically characterized by the apparent loss of a nucleosome and the generation of a nucleosome-free region within a gene’s promoter (1). However, nucleosome sliding can also change the availability of regulatory sequences as we have recently described in Simian Virus 40 (SV40) chromatin (2).

Presently, there are three procedures that are generally used to map the location of nucleosomes in chromatin; micrococcal nuclease digestion (3) and Illumina ATAC-Seq (4) are typically used to map bulk nucleosomes while ChIP-Seq (5) is used to map nucleosomes containing specific modified histones. Micrococcal nuclease digestion is used because the enzyme prefers to digest double-strand DNA when both strands at the same site are exposed. Thus, there is a clear preference for cleavage in linker DNA over DNA present in a nucleosome when this enzyme is used. Because of this relative specificity, micrococcal nuclease has been used extensively for the analysis of nucleosomes in chromatin. In order to map the location of nucleosomes by DNA sequencing using micrococcal nuclease, the DNA fragments generated by the nuclease must be purified and sequencing libraries prepared from the DNA. Nucleosomes can also be mapped using the ATAC procedure. The ATAC procedure is an in vitro transposon based strategy in which a transposase (Tagment from Illumina) targets linker DNA in regions which are open in the chromatin. The target chromatin is fragmented and transposed on to linkers which allows for the direct formation of sequencing libraries as part of the fragmentation reaction. Depending upon the nature of the target chromatin ATAC-Seq can be used either to identify open regions in total chromatin or the position of nucleosomes in the regions of chromatin susceptible to transposition. Nucleosomes containing specific histone modifications are typically mapped using ChIP-Seq. An antibody recognizing the target histone modifications is used to bind to fragmented chromatin, the bound chromatin purified for library preparation, and the resulting libraries sequenced. In each of these techniques the reads obtained from sequencing are mapped against the genome of interest to determine the location of nucleosomes.

While all three procedures have been extensively used successfully, each also has certain limitations. For example, for micrococcal nuclease the concentration of enzyme and length of digestion must be titrated for each batch of chromatin in order to obtain optimal digestion for studying nucleosomes (3). As described above ATAC-Seq is very sensitive to higher-order structure and for this reason is used primarily to investigate regions of open chromatin (4). ChIP-Seq while a very powerful technique to locate nucleosomes carrying a modified histone of interest does not generate information about nucleosome location lacking the modified histone of interest (5). Recently, New England Biolabs has made available a kit, the NEBNext Ultra II FS DNA Library Kit for Illumina, for the preparation of sequencing libraries based upon a strategy different from the three available techniques. With the NEB FS kit chromatin is fragmented using a proprietary process which cleaves naked DNA in a sequence independent manner (persona communication New England Biolabs). We report the successful use of this kit for the mapping of nucleosomes in chromatin from Simian Virus 40 (SV40) obtained at various times during the life cycle of the virus.

## Materials and Methods

### Preparation of SV40 chromatin

SV40 chromatin was prepared from BSC-1, CCL26, monkey kidney cells (ATCC) infected with SV40 strain 776. Techniques for preparing SV40 minichromosomes, virus, and chromatin from virus have been described in detail (6).

### MN-Seq

The procedures which we used to prepare sequencing libraries by MN-Seq have been previously described in detail (6, 7). For each sample of SV40, we determined the optimum extent of digestion by incubating the chromatin with diluted micrococcal nuclease in digestion buffer at 4° for from 5 to 30 seconds. The amount of dilution of the enzyme was determined for each individual preparation of SV40 chromatin. In a final volume of 50 μL SV40 chromatin (43 μL), enzyme 10X (5 μL), bovine serum albumin, 10mg/ml (1 μL) and diluted micrococcal nuclease (1 μL) were mixed. Immediately following digestion the reaction was stopped in the binding buffer from a Zymo ChIP DNA Clean and Concentrator Kit (Zymo Research, D5205) and purified following the protocol in the kit. Purified DNA was eluted in 25 μL DNAase and RNAase free water. Digestion conditions were monitored by real-time PCR. For the preparation of sequencing libraries, we only used digestions in which there was approximately a shift of three cycles in the real-time PCR from an undigested control to the digested sample. This corresponded to approximately 90% of the chromatin digested (at least at the site of PCR amplification).

### ChIP-Seq

The protocol which we have followed for the preparation of samples for ChIP-Seq has been described in general terms (2) and in detail (6). Samples were prepared using ChIP-validated antibody targeting hyperacetylated H4 (06-866, Millipore). ChIPs were performed using reagents and the protocol from the Millipore ChIP kit. Typically, 10 μL of antibody (1 ug/μL) was used for each reaction. For the ChIP using SV40 chromatin isolated 30 minutes post-infection we substituted a mixture of protein A and protein G magnetic beads (16-663, Millipore) for the protein A and protein G agarose which we previously used in our publications. In addition, antibody and chromatin was incubated at 4° for 4 hours prior to the addition of the A&G magnetic beads. Incubation was continued overnight and the chromatin bound to the magnetic beads washed as described. The bound chromatin was fragmented by sonication, washed twice with TE buffer from the kit, and eluted from the beads in the elution buffer supplied with the kit. The eluted DNA fragments were purified on a Monarch PCR & DNA Cleanup Kit (#T1030S, New England BioLabs and eluted with 26 μL DNAase and RNAase free water. The eluted samples were then lyophilized to dryness.

### ATAC-Seq

Libraries were prepared directly for ATAC-Seq using the Illumina Nextera DNA Library Prep Kit and PCR amplified using the Illumina Nextera Index Kit. Using reagents from the kit, SV40 chromatin (22.5 μL) was incubated with the 2X buffer from the kit (25 μL) and Tagment enzyme (2.5 μL) for 30 minutes at 37°. Following the fragmentation of the chromatin by enzyme the reaction, the fragmented DNA was purified using the Zymo ChIP DNA Clean and Concentrator Kit (Zymo Research, D5205). DNA was eluted from each column using 16 μL of DNAase and RNAase free water. (2, 7). Libraries were size-selected by submerged agarose gel electrophoresis. A band was excised from the gel corresponding to a size between 200 and 300 base pairs, the band was purified using a Zymo Research Gel DNA Recovery kit, and the size-selected sub-library amplified by PCR. The amplified libraries were then purified with AMPure and eluted in 16 μL of DNAase and RNAase free water.

### FS-Seq

SV40 chromatin was fragmented using the NEBNext UltraII FS DNA Library Prep Kit for Illumina (#E7805S). SV40 chromatin (13 μL) was incubated with the kit’s reaction buffer (3.5 μL) and enzyme mix (1 μL) for various times but typically 5 minutes at 4°. The fragmentation reaction was stopped by dilution of the reaction mixture in Monarch PCR & DNA Cleanup Kit (#T1030S) DNA binding buffer (100 μL). The DNA was then purified using the cleanup kit according to the protocol supplied with the kit. Fragmented DNA was eluted from the column with DNAase and RNAase free water (26 μL). An aliquot of the chromatin used for fragmentation (1 μL) and the eluted DNA (1μL) was amplified by PCR with primers which recognize the early coding region as previously described (6) in order to determine the extent of fragmentation. The time of fragmentation was adjusted so that we observed a three-cycle reduction in the threshold cycle of the amplification when comparing the fragmented aliquot to the input chromatin. Following PCR analysis, the samples were lyophilized to dryness. The purified fragmented DNA was used for library preparation as described in the next section.

### Library preparation

Libraries were prepared from purified DNA (except for ATAC-Seq) after drying using the NEB Next Ultra II Library Prep kit for Illumina (#E7645S) for the FS-Seq samples and ChIP-Seq samples following the protocol in the kit with one exception, we used half volumes of all reagents and volume of input DNA (2, 7). Libraries were prepared for MN-Seq using Illumina TruSeq reagents and protocols. Libraries for ATAC-Seq were prepared directly using the ATAC-Seq kit. Libraries were size-selected by submerged agarose gel electrophoresis. A band was excised from the gel corresponding to a size between 200 and 300 base pairs, the band was purified using a Zymo Research Gel DNA Recovery kit, and the size-selected sub-library amplified by PCR. The amplified libraries were then purified with AMPure and eluted in 16 μL of DNase and RNase free water.

### Next Generation Sequencing and Bioinformatics

Using an Agilent Bioanalyzer, libraries were first analyzed for quality and quantity, and libraries which did not meet the quality or quantity requirements were not used for sequencing. All libraries were sequenced in the sequencing core laboratory at the University of North Dakota on an Illumina MiSeq using protocols and reagents from Illumina.. For each form of SV40 chromatin and mapping technique a minimum of four separate biological replicate libraries were prepared for sequencing with the exception of the ATAC-Seq sample for which only three biological replicates were used. The actual number of replicates that were sequenced for each combination of sample and mapping technique are indicated in the figure legends. Following MiSeq sequencing the FastQ files were subjected to an initial quality control analysis using FastQC v.0.11.2 (8). The adapters attached to the 3’ end of the reads were removed using scythe v0.981 (9). Quality trimming was carried out using sickle v1.33 (10) with a phred score of 30 as the quality threshold; reads with a length less than 45 bp were discarded. Contaminating reads from African green monkey monkey (*Chlorocebus sabeus*1.1) or human (hg19) genomes were removed following alignment to their respective genomes. The remaining unmapped reads were then aligned to the SV40 genome (RefSeq Acc: NC_001669.1), cut at nucleotide (nt) 2666, using Bowtie2 v2.2.4 (11). Duplicate reads were removed using the Picard Tools (Broad) Mark Duplicates function. Bam files from each biological replicate were filtered to contain only fragments between 100-150bp or 75-99bp using an awk script. The replicate bam files were then merged together using samtools v1.3.1 (12). Bedgraphs normalized to 1x Coverage were generated from filtered, deduplicated reads using Deeptools v2.5.4 (13). Heatmaps were made using the Z-scores of the normalized coverage, and displayed using IGV v2.3.52 (14).

## Results

Since the FS kit was likely to be inhibited by the presence of proteins in chromatin, we first determined whether the enzymes could fragment SV40 chromatin. Using real-time PCR amplification of a 300 bp SV40 fragment, we measured the relative amount of DNA in chromatin samples before digestion and after digestion at different temperatures and for different times as previously described (7). In this assay, if chromatin is digested the amount of DNA in the sample is reduced and the magnitude of the reduction is an indication of how extensively the chromatin has been fragmented. Since we observed that SV40 chromatin was digested by the enzymes in the FS kit (data not shown), we then used the kit to prepare libraries for sequencing. However, because the amount of SV40 chromatin that we work with is relatively low, we found that the standard conditions recommended by the kit resulted in too much digestion. For this reason, we modified the protocol in order to optimize it for analyzing small viral chromatin like SV40. Digestions were all done at 4° C for 5 minutes and then quenched in binding buffer from the NEB Monarch PCR and DNA Cleanup Kit. The DNA in the binding buffer was then purified according to the protocol in the kit. The purified DNA was then used for the preparation of libraries using the NEB Next Ultra II Library prep kit and the libraries sequenced.

The general strategy (6) that we have been following for mapping nucleosomes on the SV40 genome has been to first prepare libraries from chromatin fragmented according to a particular procedure and to then size select the fragments containing nucleosome-sized inserts by submerged agarose gel electrophoresis on low-melting agarose. The purpose of this size-selection is to enrich for fragments that are likely to result from a mononucleosome. Typically, we select inserts between 75 and 250 bp in length. The size selected sub-fraction of the initial library is purified, PCR amplified, and sequenced. Following sequencing, as part of our bioinformatics analysis of the resulting sequencing data, we plot only those reads which are between 100 and 150 bp in length to identify nucleosomes. We focus on these reads because they are most likely to be generated by mononucleosomes. Shorter reads might be derived from linker DNA or other protein-protected complexes in the chromatin while longer reads might be derived from nucleosomes with linker DNA attached or dinucleosomes. It would be impossible to know where a nucleosome was actually located in the latter reads because of their size. Both of these types of reads would add confusion to our analysis of nucleosome location.

In order to determine whether the fragmentation of SV40 chromatin found in disrupted virions by the FS kit was consistent with the generation of nucleosome fragments, we compared the sequencing results from our FS libraries to data we have previously generated using micrococcal nuclease (7) and ChIP-Seq with antibody to hyperacetylated H4 (HH4) (2) or the Illumina ATAC-Seq kit. The latter libraries were prepared according to the protocol in the kit. The results of this comparison are shown in Figure 1 as heatmaps with the intensity of the yellow color corresponding to the frequency in which sequencing reads are found at a particular site in the SV40 genome. As shown in this figure we observed the brightest yellow bands over the enhancers found in the SV40 regulatory region using FS-Seq, ATAC-Seq, and ChIP-Seq. While there was a relatively faint band using MN-Seq which corresponded to the bright band found with the other techniques, pattern of bands was somewhat different with MN compared to all of the other procedures. To the right and left of the bright band located over the enhancer were bands corresponding to the transcription start sites for early and late transcription respectively which appeared with all of the techniques to a greater or lesser extent. In addition to these relatively brighter bands located in the major SV40 regulatory region, there were a number of other fainter bands that appeared to coincide between FS-Seq and one or more of the other techniques. In particular there were a number of bands that seemed to coincide between FS-Seq and ChIP-Seq for SV40 chromatin from disrupted virions.

**Figure 1.**
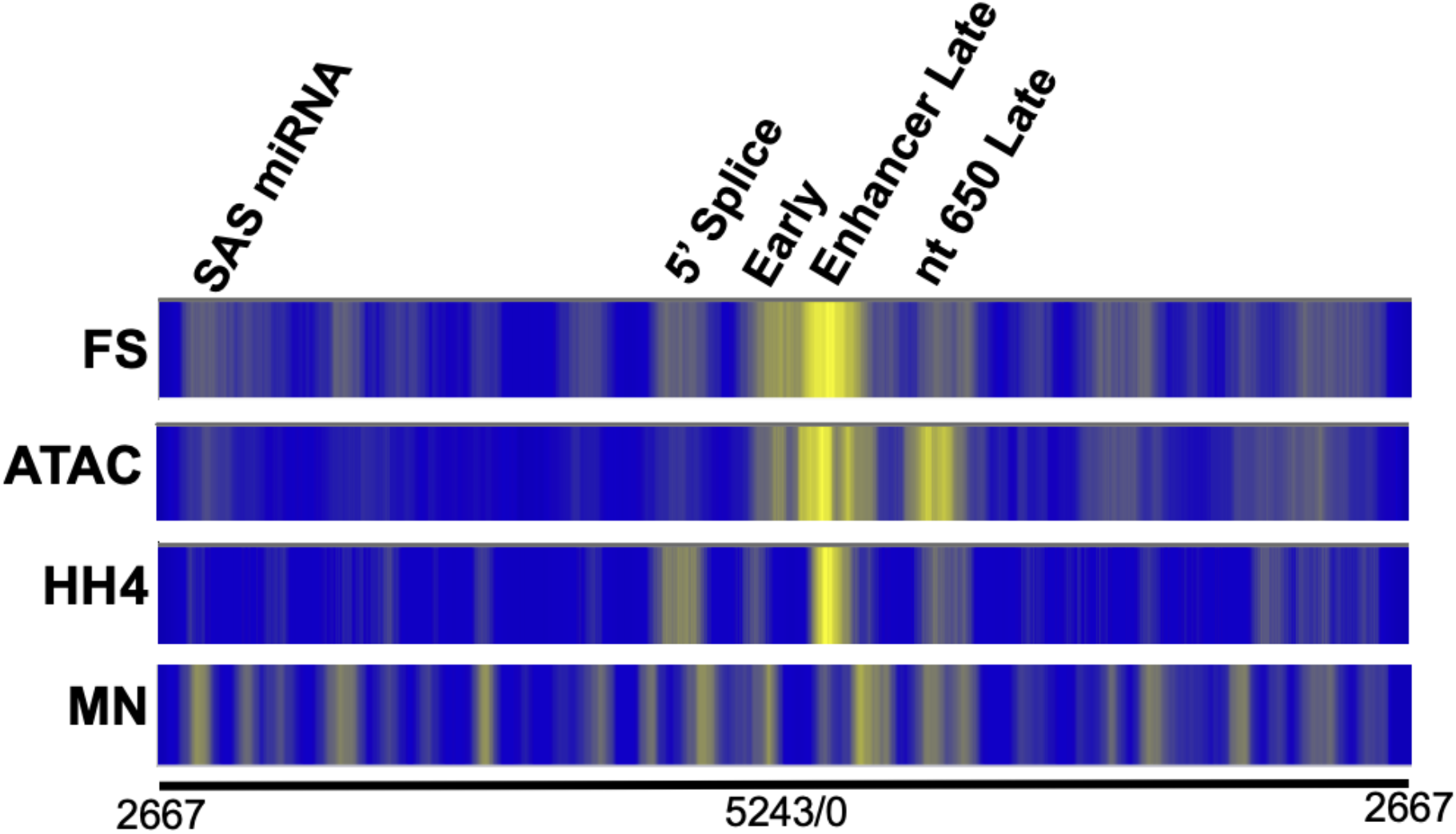
A comparison of the location of nucleosomes in chromatin from disrupted SV40 virions using FS-Seq, ATAC-Seq, ChIP-Seq, and MN-Seq. Using SV40 chromatin obtained from disrupted virions and FS-Seq, ATAC-Seq, ChIP-Seq, and MN-Seq procedures, libraries were obtained, and paired-end sequenced on an Illumina MiSeq. Heatmaps were prepared from the sequencing data (reads from 100-150 base pairs in length) from at least 4 biological replicates, normalized, merged, and plotted against the SV40 genome linearized between nt 2666 and nt2667 on the X-axis. The intensity of the yellow is a schematic representation of the number of reads obtained from a given region of the SV40 genome. FS-Seq, libraries prepared using the NEBNext UltraII FS kit (4 biological replicates), ATAC-Seq, libraries prepared with the Illumina Nextera kit (3 biological replicates), ChIP-Seq, libraries prepared from DNA present following ChIP analysis with antibody to hyperacetylated H4 (HH4) (5 biological replicates, MN-Seq, libraries prepared from chromatin digested with micrococcal nuclease (5 biological replicates).

We have previously observed that the pattern of nucleosomes on the SV40 genome changes over the course of an infection when analyzed by MN-Seq and ChIP-Seq (2, 7). In order to determine whether the pattern of nucleosome positioning when analyzed by FS-Seq also changed consistent with our previous observations, we compared results from FS-Seq to the two other techniques in SV40 chromatin isolated at 30 minutes post-infection and 48 hours postinfection. In our previous work with MN-Seq (7) and ChIP-Seq (2) we observed using MN-Seq that a nucleosome was present in the enhancer in chromatin from 48 hour infections which appeared to be present in much less of the chromatin in virions or at 30 minutes post infection. Using ChIP-Seq we observed a sliding of the enhancer nucleosome during the process of encapsidation of SV40 chromatin late in infection (2). In this analysis we also included ChIP-Seq data showing the location of RNA Polymerase II (RNAPII),

The results of this comparison are shown in Figure 2. We observed that many of the brighter bands in the heatmap from FS-Seq using chromatin from virions, were also observed in the SV40 chromatin isolated at 30 minutes and 48 hours post-infection although with different intensities. This result suggests that FS-Seq was fragmenting SV40 chromatin in a very consistent way. The differences in intensity suggested that fragmentation was also sensitive to differences in the specific chromatin structure present in the SV40 samples depending upon the source. The changes in intensity of the bands was particularly noticeable at the early and late transcription starts, with a major increase at the early start late in infection and an increase at the late start early in infection. These changes would be consistent with nucleosome positioning acting as a competitor for binding by RNA Polymerase II.

**Figure 2.**
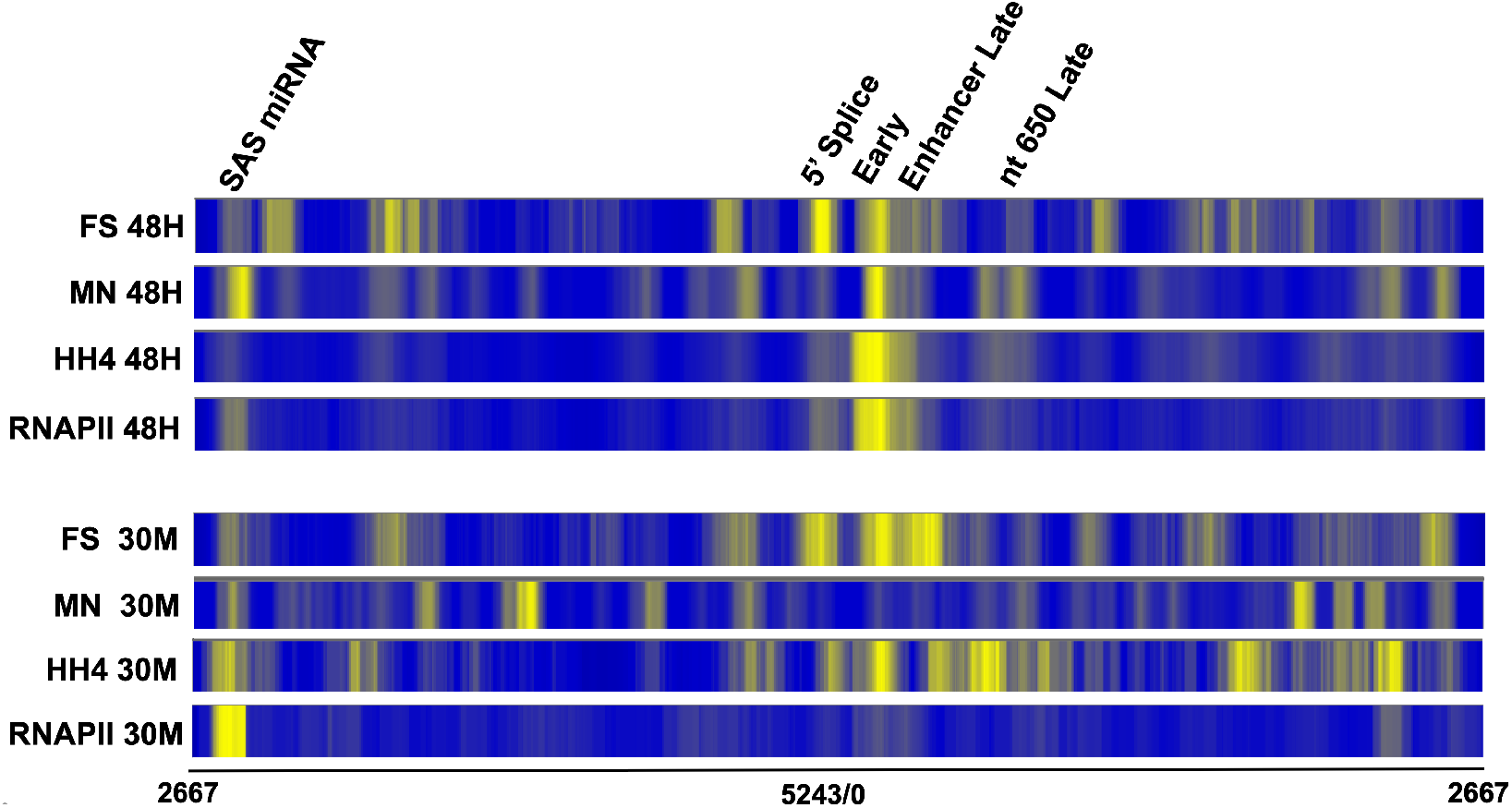
A comparison of the location of nucleosomes in SV40 chromatin from minichromosomes isolated 30 minutes and 48 hours post-infection using FS-Seq, ChIP-Seq, and MN-Seq. SV40 chromatin was obtained from minichromosomes isolated from cells infected for either 30 minutes or 48 hours with SV40. Libraries were prepared using either FS-Seq, ChIP-Seq (anti-HH4), or MN-Seq procedures and paired-end sequenced on an Illumina MiSeq. Heatmaps were prepared from the sequencing data (reads from 100-150 base pairs in length) from at least 4 biological replicates, normalized, merged, and plotted against the SV40 genome linearized between nt 2666 and nt2667 on the X-axis. The intensity of the yellow is a schematic representation of the number of reads obtained from a given region of the SV40 genome. FS-Seq, libraries prepared using the NEBNext UltraII FS kit (30 min, 7 biological replicates, 48 hours, 4 biological replicates), ChIP-Seq, libraries prepared from DNA present following ChIP analysis with antibody to hyperacetylated H4 (HH4) (30 min, 4 biological replicates, 48 hours, 7 biological replicates), or RNAPII (30 mins 5 biological replicates, 48 hours, 8 biological replicates), MN-Seq, libraries prepared from chromatin digested with micrococcal nuclease (30 min, 5 biological replicates, 48 hours, 4 biological replicates). FS indicates samples prepared using the FS kit. HH4 indicates samples were prepared from ChIPs with HH4. MN indicates samples were prepared using micrococcal nuclease. 48H indicates chromatin from SV40 minichromosomes isolated 48 hours post-infection. 30M indicates chromatin from SV40 minichromosomes isolated 30 minutes post-infection.

Perhaps, not surprisingly, we observed some similarities as well as some significant differences in comparisons between FS-Seq, MN-Seq, and ChIP-Seq with antibody to HH4. Comparing FS-Seq to ChIP-Seq we observed many of the brighter bands located at the same place by both procedures. This was true for the enhancer band and other bands in the major regulatory region in all forms of SV40 chromatin. We also observed overlaps between many of the other bands in the early and late coding regions. However, there were also bands present in the heatmaps from FS-Seq or ChIP-Seq which did not seem to be present in the other preparation. A similar type of result was also observed in comparison to MN-Seq with one major difference. In the MN-Seq the bands present in the regulatory region in chromatin from virions and 30-minute infections were much lighter than the corresponding bands from FS-Seq and ChIP-Seq. This result suggests that MN-Seq was preferentially digesting the chromatin located in this region, while FS-Seq and ChIP-Seq did not.

While our primary purpose in these studies was to determine whether FS-Seq was a viable alternative to MN-Seq or ATAC-Seq, we also tested whether FS-Seq might be useful for identifying the location of certain transcription factors. In the SV40 genome the transcription factor SP1 potentially can bind to a region that is approximately 60 base pairs in length that is located between nucleotide 40 and nucleotide 104 (15–17). Because SP1 binding to its target DNA is relatively strong, we hypothesized that it might protect its target DNA from fragmentation by the FS kit. However, because the fragment of SV40 DNA protected by SP1 would be expected to be much smaller than the size that we typically use for analyzing nucleosome location, 100-150 bp, for the analysis of transcription factor binding we used only the reads from 75 to 99 bp in size.

The result of this analysis using SV40 chromatin isolated 48 hours post infection is shown in Figure 3 in which we show a comparison between the location of RNAPII by ChIP-Seq using reads from 100-150 bp in length, the location of SP1 by ChIP-Seq using only the reads from 75-99 bp in length, the location of nucleosomes by FS-Seq using the reads from 100-150 bp in length, and the location of protected DNA fragments by FS-Seq using only the reads from 75-99 bp in length. As is apparent from the figure the pattern of reads for the short fragments by FS-Seq is very different from what is obtained using nucleosome-sized sequencing reads. With the shorter reads there appears to be only a single major band which corresponds very well to the location of the major SP1 band in the regulatory region found by ChIP-Seq and interestingly to the location of RNAPII as well. These results suggest that FS-Seq combined with a bioinformatics analysis focused on shorter sequencing reads may be useful to identify the location of RNAPII or transcription factors which bind tightly to DNA and protect relatively large regions of DNA.

**Figure 3.**
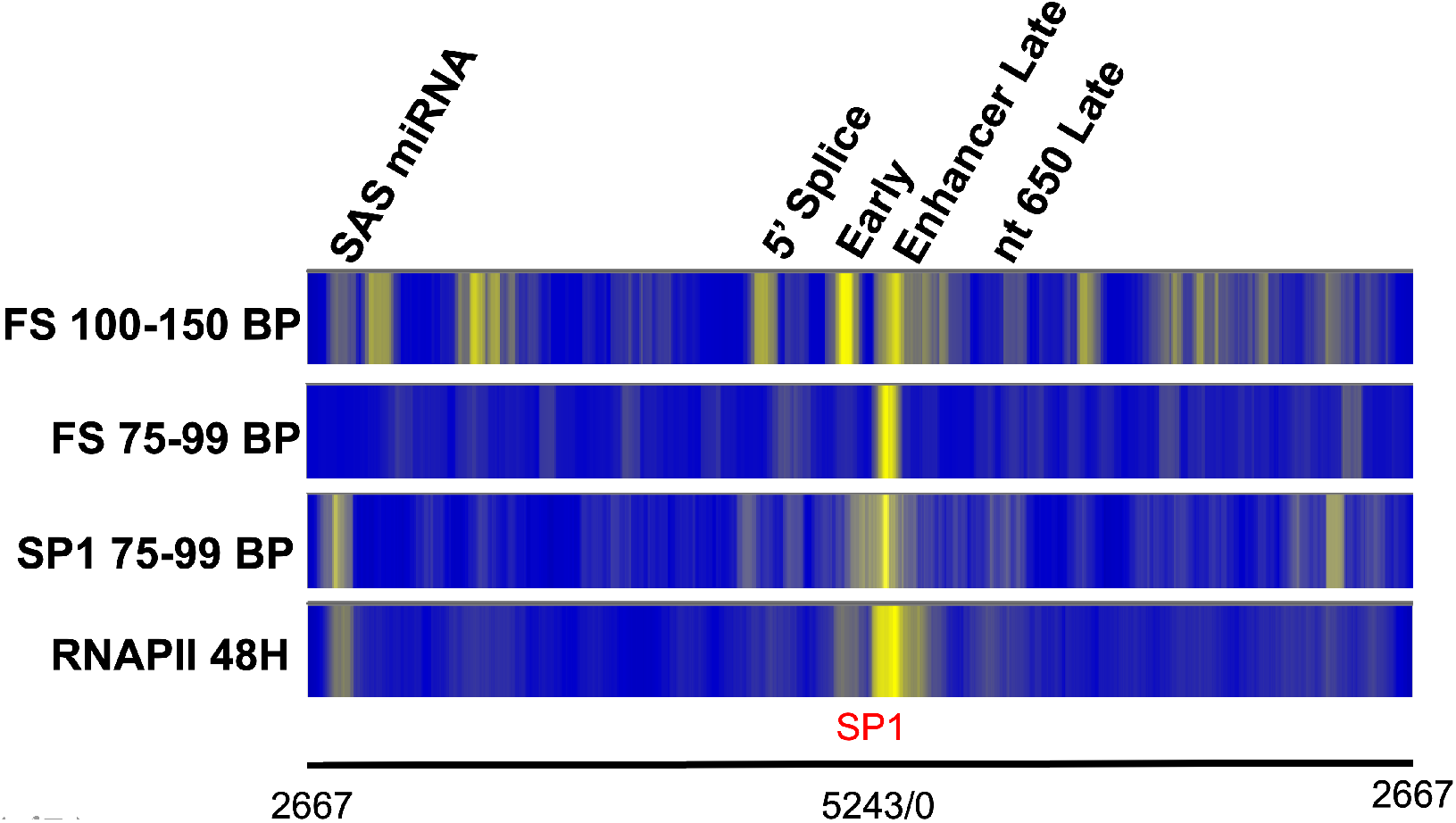
A comparison of the location of SP1, RNAPII, and the location of sequencing reads from 100-150 base pairs in length and 75-99 base pairs in length obtained by FS-Seq from SV40 chromatin obtained 48 hours post infection. SV40 chromatin was obtained from minichromosomes isolated from cells infected with SV40 for 48 hours. Libraries were prepared using either FS-Seq, or ChIP-Seq (anti-RNAPII or SP1) and paired-end sequenced on an Illumina MiSeq. Heatmaps were prepared from the sequencing data from at least 4 biological replicates, normalized, merged, and plotted against the SV40 genome linearized between nt 2666 and nt2667 on the X-axis. The size of the merged reads is indicated in the figure. Reads from 100-150 base pairs in length are typically used to identify the location of nucleosomes. Reads from 75-99 base pairs in length would be expected to reflect protection of smaller fragments of chromatin. The intensity of the yellow is a schematic representation of the number of reads obtained from a given region of the SV40 genome. FS-Seq, libraries prepared using the NEBNext UltraII FS kit, 4 biological replicates, ChIP-Seq, libraries prepared from DNA present following ChIP analysis with antibody to RNAPII, (8 biological replicates), or SP1 (5 biological replicates). FS indicates samples prepared using the FS kit. RNAPII indicates samples were prepared from ChIPs with antibody to RNA Polymerase II. SP1 indicates samples were prepared form ChIPs with antibody to SP1.

## Discussion

According to documentation from New England Biolabs, the FS kit was designed for sequence-independent fragmentation of purified genomic DNA followed by the preparation of sequencing libraries in a continuous work-flow (New England biolabs technical note “High-Yield, Scalable Library Preparation with the NEBNext UltraII FS DNA Library Prep Kit,” 2017). Since the process used for DNA fragmentation should function on linker DNA in chromatin similarly to the way that they function on naked DNA, we tested whether the kit could also be used for mapping nucleosomes using chromatin instead of DNA as the input for fragmentation. For this analysis we used SV40 chromatin obtained at different times in the virus life cycle, because we had experience analyzing SV40 chromatin structure using other techniques. In addition, we have observed certain changes in SV40 chromatin structure in different forms of SV40 chromatin and could compare these previously observed changes to what was found using the FS kit.

In order to use the FS kit for mapping nucleosomes in SV40, we found that it was necessary to do the fragmentation at 4° and for relatively short periods of time, on the order of five minutes. This was most likely due to the fact that the amount of SV40 chromatin in our samples was relatively low compared to what can be obtained from a cell. Since the amount of fragmentation will increase during the temperature shift which is done to prepare libraries in a continuous fashion, we also found it necessary to purify the fragmented DNA immediately following the low-temperature incubation. We then prepared libraries using the regular NEB Next Ultra II kit and size selected for nucleosome-sized or smaller fragments in our libraries. Using this strategy, we were able to obtain sequencing libraries consistent with those which we have previously obtained. For consistency with the names for other sequencing techniques, we refer to sequencing using the FS kit as FS-Seq.

In comparing FS-Seq to MN-Seq, ChIP-Seq, and ATAC-Seq using the same samples of chromatin from disrupted SV40 virions, we found that each of the bands in its heatmap appeared to be present in at least one of the heatmaps from another sequencing technique. With the exception of the nucleosome band in the enhancer, FS-Seq appears to be most similar to the results that we previously obtained with MN-Seq although there are differences in the relative intensities of the bands in the two heatmaps. Within the SV40 regulatory region FS-Seq is very similar to the results obtained from ChIP-Seq and ATAC-Seq. Not surprisingly each of the sequencing techniques also yielded sufficiently different results suggesting that they were targeting chromatin in different ways.

For example, the major nucleosome band in the heatmaps within the enhancer was dramatically reduced using MN-Seq. Based upon how micrococcal nuclease digestion occurs there are two likely reasons for the apparent relative reduction in the presence of a nucleosome in the enhancer region using this technique. MN digestion can result in smaller than nucleosome sized fragments if digestion is carried out for longer periods of time or at higher temperatures (3). We have previously noted that this was a problem in our publication (7) because we are working with relatively small amounts of chromatin when using SV40. Secondly, MN digestion is sensitive to both AT rich regions and to what are operationally referred to as “fragile nucleosomes” (3). Fragile nucleosomes are those that are typically found in regulatory regions that are particularly sensitive to MN digestion (3).

ChIP-Seq would be expected to contain fewer bands corresponding to nucleosome positions in the heatmaps because only those nucleosomes which contain the target histone modification would be expected to be present (5). ChIP-Seq can also be subject to genomic bias since sonication and nuclease digestion potentially can occur preferentially at regions of open chromatin (5). This may account for the intensity of the band corresponding to a nucleosome in the SV40 regulatory region since open chromatin is found adjacent to this site (18–21).

ATAC-Seq might also be expected to contain fewer bands than FS-Seq with the bands present being located primarily in the regulatory region (4). Although originally developed as an alternative or complement to MN-Seq, ATAC-Seq is now generally used as a way to identify open chromatin (4). Since SV40 chromatin typically has an open region associated with the regulatory region, it is not surprising that the brightest bands in the heatmap are found in the regulatory region using ATAC-Seq with very few bands in the coding regions.

In many respects FS-Seq using SV40 virion chromatin appears to resemble a mixture of the results from MN-Seq and ATAC-Seq. The brightest bands in the heatmap are located in the regulatory region like ATAC-Seq but there are also a number of other bands located at similar positions to what we obtained previously with MN-Seq. We conclude that FS-Seq like the other techniques is likely to preferentially target nucleosomes present in open regions although it can also target other accessible nucleosomes like MN-Seq.

In our previous studies on SV40 chromatin structure using MN-Seq and ChIP-Seq, we have analyzed the SV40 chromatin obtained at different times in infection or from virions in order to determine whether the location of nucleosomes remained constant or changed during the course of an infection. The former would suggest that the technique was locating nucleosomes consistently, while the latter would indicate that changes were occurring as a consequence of the viral life cycle. The results with MN-Seq and ChIP-Seq both indicated that changes occurred in the major regulatory region but tended to be consistent in other regions of the genome. The results with FS-Seq were also similar in this regard. We observed significant differences in intensity of bands in the regulatory region with FS-Seq. Notably at 48 hours post-infection the brightest band was present over the early transcription start consistent with a potential role in blocking early transcription at late times. Although bands were also found at the early start site early in infection (30 minutes), the most prominent bands were located in the enhancer and at the late transcription start suggesting that a nucleosome may be blocking the late transcription start. Each of the three mapping procedures show definite differences in the chromatin structure of the regulatory region, although with the possible exception of the sliding enhancer nucleosome seen with ChIP-Seq and to a lesser extent with FS-Seq the differences are not the same from technique to technique.

Since each mapping technique generates data which can be both consistent with and different from other mapping techniques depending upon the location of a nucleosome, we believe that combining FS-Seq with other mapping techniques would be a useful strategy to confirm nucleosome positions in chromatin of interest. FS-Seq is particularly suited as a complementary mapping technique for nucleosomes because it is technically very easy to do, gives relatively robust results, and is cheaper than other techniques.

Our results using shorter reads with FS-Seq suggest that FS-Seq may also be useful to identify potential transcription factor binding sites. A number of techniques have been utilized for this purpose, but they typically depend upon knowing the transcription factor of interest and use ChIP-Seq or comparable procedure to identify the location of the factor of interest in chromatin. The FS-Seq strategy described here does not depend upon knowing the nature of the fragment but whether the strategy will be generally useful as a global strategy to identify factor binding sites will depend upon further studies.

## Acknowledgements

The authors would like to thank Hannah Ness and the Epigenetic Core Laboratory at the University of North Dakota for sequencing the samples and the bioinformatics analyses. This work was funded by grants from the National Institutes of Health, AI094441 (to B.M.), AI142011 (to BM), GM104360 (to UND Epigenetics Core), and GM128729 (to Dakota Cancer Collaborative on Translational Activity).

## References

1. Lorch Y, Kornberg RD. 2017. Chromatin-remodeling for transcription. Q Rev Biophys 50:e5.

2. Kumar MA, Kasti K, Balakrishnan L, Milavetz B. 2018. Directed Nucleosome Sliding in SV40 Minichromosomes During the Formation of the Virus Particle Exposes DNA Sequences Required for Early Transcription. J Virol doi:10.1128/JVI.01678-18.

3. Voong LN, Xi L, Wang JP, Wang X. 2017. Genome-wide Mapping of the Nucleosome Landscape by Micrococcal Nuclease and Chemical Mapping. Trends Genet 33:495–507.

4. Sun Y, Miao N, Sun T. 2019. Detect accessible chromatin using ATAC-sequencing, from principle to applications. Hereditas 156:29.

5. Park PJ. 2009. ChIP-seq: advantages and challenges of a maturing technology. Nat Rev Genet 10:669–680.

6. Balakrishnan L, Milavetz B. 2017. Epigenetic Analysis of SV40 Minichromosomes. Curr Protoc Microbiol 46:14F 13 11–14F 13 26.

7. Kumar MA, Christensen K, Woods B, Dettlaff A, Perley D, Scheidegger A, Balakrishnan L, Milavetz B. 2017. Nucleosome positioning in the regulatory region of SV40 chromatin correlates with the activation and repression of early and late transcription during infection. Virology 503:62–69.

8. Andrews S. 2010. FastQC: a quality control for high throughput sequence data.

9. Buffalo V. 2011. Scythe: A bayesian adapter trimmer.

10. Joshi NA, Fass JN. 2011. sickle - A windowed adaptive trimming tool for FASTQ files using quality.

11. Langmead B, Salzberg SL. 2012. Fast gapped-read alignment with Bowtie 2. Nat Methods 9:357–359.

12. Li H. HB, Wysoker A., Fennell T., Ruan J., Homer N., Marth G., Abecasis G., Durbin R. and 1000 Genome Project Data Processing Subgroup. 2009. The Sequence alignment/map (SAM) format and SAMtools. Bioinformatics 25:2078–2079.

13. Ramirez F, Ryan DP, Gruning B, Bhardwaj V, Kilpert F, Richter AS, Heyne S, Dundar F, Manke T. 2016. deepTools2: a next generation web server for deep-sequencing data analysis. Nucleic Acids Res 44:W160–165.

14. Robinson JT, Thorvaldsdottir H, Winckler W, Guttman M, Lander ES, Getz G, Mesirov JP. 2011. Integrative genomics viewer. Nat Biotechnol 29:24–26.

15. Dynan WS, Tjian R. 1983. Isolation of transcription factors that discriminate between different promoters recognized by RNA polymerase II. Cell 32:669–680.

16. Dynan WS, Tjian R. 1983. The promoter-specific transcription factor Sp1 binds to upstream sequences in the SV40 early promoter. Cell 35:79–87.

17. Barrera-Saldana H, Takahashi K, Vigneron M, Wildeman A, Davidson I, Chambon P. 1985. All six GC-motifs of the SV40 early upstream element contribute to promoter activity in vivo and in vitro. Embo J 4:3839–3849.

18. Kondoleon SK, Kurkinen NA, Hallick LM. 1989. The SV40 nucleosome-free region is detected throughout the virus life cycle. Virology 173:129–135.

19. Scott WA, Wigmore DJ. 1978. Sites in simian virus 40 chromatin which are preferentially cleaved by endonucleases. Cell 15:1511–1518.

20. Varshavsky AJ, Sundin OH, Bohn MJ. 1978. SV40 viral minichromosome: preferential exposure of the origin of replication as probed by restriction endonucleases. Nucleic Acids Res 5:3469–3477.

21. Kube D, Milavetz B. 1989. Generation of a nucleosome-free promoter region in SV40 does not require T-antigen binding to site I. Virology 172:100–105.

